# Large-scale transgenic *Drosophila* resource collections for loss- and gain-of-function studies

**DOI:** 10.1101/852376

**Authors:** Jonathan Zirin, Yanhui Hu, Luping Liu, Donghui Yang-Zhou, Ryan Colbeth, Dong Yan, Ben Ewen-Campen, Rong Tao, Eric Vogt, Sara VanNest, Cooper Cavers, Christians Villalta, Aram Comjean, Jin Sun, Xia Wang, Yu Jia, Ruibao Zhu, Pin Peng, Jinchao Yu, Da Shen, Yuhao Qiu, Limmond Ayisi, Henna Ragoowansi, Ethan Fenton, Senait Efrem, Annette Parks, Kuniaki Saito, Shu Kondo, Liz Perkins, Stephanie E. Mohr, Jianquan Ni, Norbert Perrimon

**Affiliations:** Department of Genetics, Blavatnik Institute, Harvard Medical School, Boston, Massachusetts 02115; Shanghai Institute of Plant Physiology and Ecology, Chinese Academy of Sciences, 200032 Shanghai, China; Tsinghua University, Tsinghua Fly Center, Beijing, 100084, China; Bloomington *Drosophila* Stock Center, Bloomington, Indiana, 47405; Invertebrate Genetics Laboratory, National Institute of Genetics, Mishima, Shizuoka, 411-8540, Japan; Howard Hughes Medical Institute, Boston, Massachusetts 02115

**Keywords:** Drosophila, RNAi, CRISPR, Cas9, knockout, overexpression, phenotypes, screens

## Abstract

The Transgenic RNAi Project (TRiP), a *Drosophila* functional genomics platform at Harvard Medical School, was initiated in 2008 to generate and distribute a genome-scale collection of RNAi fly stocks. To date, the TRiP has generated >15,000 RNAi fly stocks. As this covers most *Drosophila* genes, we have largely transitioned to development of new resources based on CRISPR technology. Here, we present an update on our libraries of publicly available RNAi and CRISPR fly stocks, and focus on the TRiP-CRISPR overexpression (TRiP-OE) and TRiP-CRISPR knockout (TRiP-KO) collections. TRiP-OE stocks express sgRNAs targeting upstream of a gene transcription start site. Gene activation is triggered by co-expression of catalytically dead Cas9 (dCas9) fused to an activator domain, either VP64-p65-Rta (VPR) or Synergistic Activation Mediator (SAM). TRiP-KO stocks express one or two sgRNAs targeting the coding sequence of a gene or genes, allowing for generation of indels in both germline and somatic tissue. To date, we have generated more than 5,000 CRISPR-OE or -KO stocks for the community. These resources provide versatile, transformative tools for gene activation, gene repression, and genome engineering.

## INTRODUCTION

*Drosophila* is an exemplary model for genetic studies and has been used for decades to identify genes involved in developmental and cellular processes [reviews include (Nagy et al. 2003; Venken and Bellen 2005; McGurk et al. 2015; Ugur et al. 2016; Sonoshita and Cagan 2017)]. Strengths of the *Drosophila* model include (1) the extraordinary evolutionary conservation between insects and vertebrates of many of the basic processes of development and nearly all of the basic signal transduction mechanisms and transcriptional regulators and (2) the suite of sophisticated technologies available for manipulating the *Drosophila* genome and for selection and analysis of mutant phenotypes. The relevance of *Drosophila* to humans is best illustrated by the observations that >60% of human genes associated with diseases have counterparts in *Drosophila* (Rubin et al. 2000; Hu et al. 2011) and at least 135 human genes have been shown experimentally to rescue loss-of-function mutation of their counterparts in *Drosophila* (Fernandez-Hernandez et al. 2016). Moreover, *Drosophila* is increasingly being used to model a wide range of human diseases, from developmental to metabolic to age-related diseases and more. Notable areas in which *Drosophila* models are having an impact include a wide variety of neurological diseases [reviews include (Okray and Hassan 2013; Coll-Tane et al. 2019)]; cancer (Sonoshita and Cagan 2017); metabolic disease and diabetes (Park et al. 2014); responses to infection by human pathogens (Apidianakis and Rahme 2011); immune disorders (Bergman et al. 2017); heart disease (Viswanathan et al. 2014); and inherited disorders (Wangler et al. 2017; Mishra-Gorur et al. 2019). *Drosophila* is also increasingly being used to help identify causative variants associated with previously undiagnosed genetic disorders (Wangler et al. 2017; Splinter et al. 2018). Despite the impressive set of tools currently available for *Drosophila* genetic studies, locating and/or generating disease models in *Drosophila* can be time consuming, and at the start of our project, well-characterized loss-of-function alleles or validated RNAi and/or sgRNA strains were not available. This is a particularly limiting step for scientists who are not expert at working with *Drosophila* but have an interest in using this powerful system to gain insight into the biological functions of their genes of interest.

Genetic analysis of mutants relies on two complementary approaches, loss-of-function (LOF) and gain-of-function (GOF) studies. In LOF studies, the role of a gene is inferred from the phenotype that results from partial or complete absence of the gene product. LOF was classically achieved *in vivo* in *Drosophila* using random mutagenesis (St Johnston 2002) and in more recent years, is achieved by knockdown using RNA interference (RNAi) (Dietzl et al. 2007), or knockout using CRISPR-Cas9 (Port et al. 2014). Transgenic RNAi is a powerful and straightforward method for analysis of gene function [reviews include (Perrimon et al. 2010; Heigwer et al. 2018)]. In *Drosophila*, RNAi is cell-autonomous, and is used in conjunction with the GAL4/UAS system (Brand and Perrimon 1993) for both spatial and temporal knockdown, providing a powerful approach to the study of genes with pleiotropic functions. CRIPSR/Cas9 can also be used to generate tissue-specific mutations *in vivo* (Port et al. 2014; Port and Bullock 2016), as CRISPR/Cas9 efficiently generates double strand breaks (DSBs) that can be used to generate mutations (Bassett et al. 2013; Gratz et al. 2013; Kondo and Ueda 2013; Ren et al. 2013; Yu et al. 2013; Sebo et al. 2014; Lee et al. 2018).

Although powerful, LOF analysis alone is not sufficient to fully annotate gene functions, for example due to the lack of detectable phenotypes for many genes (Rorth et al. 1998). In particular, it has been estimated that up to 75% of *Drosophila* genes are phenotypically silent upon LOF because of genetic redundancy (Miklos and Rubin 1996). To complement LOF studies, GOF studies that use controlled mis- or over-expression of a gene may help elucidate function. In addition, GOF approaches have the potential to provide insights into gene function even in cases of redundancy and there are many examples of *Drosophila* genes which when overexpressed are associated with phenotypes such as patterning defects, aberrant cell proliferation, neurodegeneration, metabolism abnormalities, etc. Importantly, GOF analyses can also provide insight into diseases associated with mis- or over-expression. For example, genes identified as up-regulated in cancer can be tested for oncogenic activity using overexpression studies (Croce 2008). In addition, *Drosophila* has been used to help uncover specific genes associated with chromosomal duplication disorders (Grossman et al. 2011).

The available reagents for overexpression experiments using GAL4/UAS belong to two categories: (1) random insertions of an element that brings UAS upstream of a specific gene; and (2) transgenic insertions of a UAS sequence fused to a specific open reading frame (ORF). Regarding the insertion lines, many stock collections are available from the Bloomington Drosophila Stock Center (BDSC); namely, the EP (Rorth 1996), EPg (Staudt et al. 2005), EPgy2 (Bellen et al. 2004), and Exelixis (Thibault et al. 2004) collections. These have the potential to misexpress genes that flank the insertion sites in the presence of GAL4, depending on the location and orientation of the insertion. However, only a small number have proven useful to identify overexpression phenotypes as these collections have not been validated systematically to identify which insertions result in overexpression of the nearby gene(s) or to assess other effects of the insertions. To overcome this shortcoming, the Zurich ORFeome project generated fly stocks – whereby, the ORF is cloned into a UAS vector, and integrated into the genome using ΦC31 integrase – that cover about 2,850 genes, mostly transcription factors (Bischof et al. 2013; Schertel et al. 2015). However, these resources are limited in scope.

Since its inception in 2008, the Transgenic RNAi Project (TRiP) has generated reagents for *in vivo* studies in *Drosophila*. The vectors used for RNAi reagents, approach and production platform of the lines were described extensively previously (Perkins et al. 2015). Here, we describe an update of the project and focus on novel resources based on CRISPR-Cas9 produced using the same efficient platform. Specifically, we describe (1) currently available reagents for *in vivo* LOF and GOF studies based on RNAi and CRISPR technologies; (2) large-scale collections of transgenic RNAi fly stocks and more recently, transgenic sgRNA fly stocks for either overexpression or knockout, in particular for studies of orthologs of human disease-associated genes; (3) ‘toolkit’ fly stocks and reagents that facilitate optimal use of RNAi and CRISPR sgRNA fly stock resources; and (4) a database of information on the quality of individual RNAi and sgRNA lines, RSVP Plus (http://fgr.hms.harvard.edu/rsvp).

## MATERIALS AND METHODS

### Plasmid cloning

shRNAs for the transgenic RNAi stocks were cloned into the VALIUM series and pNP vectors (see Supplementary Materials). sgRNAs for the TRiP-CRISPR stocks were cloned into *pCFD3* (PORT *et al.* 2014), *pCFD4* (PORT *et al.* 2014), *pCFD6* (PORT *et al.* 2016), U6B-sgRNA2.0 (Jia *et al.* 2018), and flySAM2.0 (Jia *et al*. 2018).

### Fly genetics and embryo injections

Flies were maintained on standard fly food at 25°C, unless otherwise noted. Fly stocks were obtained from the Perrimon lab collection or BDSC (indicated with BL#). Stocks used in this study are: *y v; attP40* (BL36304); *y v; attP2* (BL36303); *y v nos-phiC31-int; attP40* (BL25709); *y v nos-phiC31-int; attP2* (BL25710); *y sc v; Dr e/TM3, Sb* (BL32261); *y sc v; Gla/CyO* (35781); *y w; Dmef2-GAL4* (Perrimon); y *w; elav-GAL4* (Perrimon); *w; ey-GAL4* (BL5534); *y w; nub-GAL4* (Perrimon); *y w; act5C-GAL4* (Perrimon); *w; myo1A-GAL4* (Perrimon); *w; Lpp-GAL4* (Perrimon); *w; Cg-GAL4* (Perrimon); *y w; tub-GAL4* (Perrimon); *w; da-GAL4* (Perrimon); *PG142-GAL4* (Perrimon); *hml-GAL4* (Perrimon); *w; UAS-dCas9.VPR; tub-GAL4TM6* (BL67065); *w; nub-GAL4; UAS-dCas9.VPR* (BL67055); *y w; CG6767[MI09551-GFSTF.2]* (BL59305); *y sc v; TOE.GS01642 = TRiP-OE CG6767* (BL81664); *y sc v; TOE.GS00152 = TRiP-OE spz* (BL67551); *y sc v; TOE.GS00045 = TRiP-OE grk* (BL 67525); *TOE.GS00016 = TRiP-OE spz4* (BL 67519); *y sc v; TOE.GS00088 = TRiP-OE spz6* (BL 67538); *y sc v; TOE.GS00062 = TRiP-OE trk* (BL 67532); *y sc v; TOE.GS01583 = TRiP-OE su(r)* (BL79460); *y sc v; SAM.dCas9.TH12432.S* = control guide (Perrimon); *y sc v*; *SAM.dCas9.TH12427.S* = control guide (Perrimon).

For embryo injections, each plasmid was column purified (Qiagen), eluted in injection buffer (100 μM NaPO4, 5 mM KCl), and adjusted to 50 ng/μl. Plasmids were injected individually or as pools into *y v nos-phiC31-int; attP40* (for chromosome 2 insertions) or *y v nos-phiC31-int; attP2* (for chromosome 3 insertions). Injected G0 flies were crossed with *y sc v; Gla/Cyo or y sc v; Dr e/TM3, Sb* to identify transformants and remove the integrase from the X-chromosome.

### flySAM lethality

Virgin females of each GAL4 line were crossed to balanced male control stocks or male flySAM2.0 sgRNA stocks. Vials were stored at 25°C and flipped every 3 days. The proportion of GAL4+sgRNA to GAL4+Bal were recorded in the resulting progeny (1 = complete viability, 0 =complete lethality). These scores were normalized to the results of the *y v; attP2* cross.

### Human Disease (HuDis) library screen

∼1200 RNAi fly stocks corresponding to 670 high confidence *Drosophila* orthologs of human disease genes were screened with ubiquitous and tissue-specific drivers. Female GAL4 lines were crossed to male RNAi lines at 25°C. Progeny were scored for lethality and visible phenotypes. Stocks were sub-categorized as lethal or viable with ubiquitous drivers. Lines for which ubiquitous expression resulted in lethality were then crossed to a panel of tissue-specific GAL4 drivers and the resulting viability and/or gross morphological phenotypes were recorded. Triglyceride and glucose levels were measured as described below for stocks that were viable with ubiquitous drivers.

### Triglyceride, Glucose and Glycogen Measurement

Six adult male flies (three biological replicates) were homogenized in 200 μL of PBS containing 0.1% Triton-X, heated at 70°C for 10 min, and centrifuged at 13,000 g for 10 min. 10 μL of supernatant was used to measure triglyceride by using Serum TG determination kits (Sigma, TR0100-1KT). 10 μL supernatant was treated with 0.2 μL water or trehalase (Megazyme, E-TREH) to digest trehalose into glucose or 1 µL amyloglucosidase (Sigma, for glycogen measurement) at 37°C for 20 min, and glucose was measured by incubation with 150 μL D-Glucose Assay reagent (Megazyme, K-GLUC) at 37°C for 5 min. Protein amounts were measured using Bradford Reagent (Sigma). Triglyceride, glucose and glycogen levels were normalized to protein amount (three technical replicates).

### qPCR

Flies of the genotype *w; UAS:dCas9-VPR; tub-Gal4/SM5, TM6B* were crossed to homozygous sgRNA lines, and the F1 were maintained for 5 d at 18°C to reduce Gal4 activity and thereby lethality. Larvae were then transferred to 27°C for 48 h, and non-Tb larvae were used for qPCR. Three to six L3 larvae were flash-frozen on dry ice and homogenized in TRIzol (Thermo Fisher); then RNA was extracted following the manufacturer’s protocol. RNA was purified using either RNeasy (Qiagen) or Direct-zol (Zymo) kits, including a 15–20 min DNase treatment, and then was used as template for first-strand synthesis and qPCR. Target genes were compared with the geometric mean of two reference genes (*Rp49* and *GAPDH*), and an annealing temperature of 57°C was used. Primers were designed using a precomputed database of *Drosophila* qPCR primers (Hu et al. 2013). All experiments were conducted in biological duplicates, with technical triplicate qPCR reactions.

## RESULTS AND DISCUSSION

### The TRiP RNAi collection

Since the description in 2015 of TRiP RNAi fly stock production and resources (Perkins et al. 2015) (Fig. 1 and Supplementary Materials), we have continued to optimize the approach and produce additional new shRNA stocks. To date, the TRiP has generated ∼15,800 RNAi stocks covering ∼9,800 unique FBgns, or 70% of the protein coding genes in the fly genome (FlyBase Release 6.30) with 85% coverage of highly conserved genes. Statistics on TRiP reagent gene coverage, vectors, target regions are shown in Table S1. TRiP lines are regularly used by *Drosophila* researchers worldwide. Currently, BDSC distributes 13,541 RNAi stocks, and to date have shipped over 700,000 TRiP stocks (Summarized in Fig. 1). In 2018 alone, BDSC sent 78,370 subcultures of TRiP stocks to 1,406 different user groups in 45 US states and 47 countries. Additional TRiP shRNA stocks are also available from the National Institute of Genetics, Japan (https://shigen.nig.ac.jp/fly/nigfly/) and the Tsinghua Fly Center, China (http://fly.redbux.cn/). Notably, the latter collection includes 1093 shRNA lines made with the new pNP vector, which functions in both in soma and germline with significantly reduced basal hairpin expression (Qiao et al. 2018).

**Figure 1:**
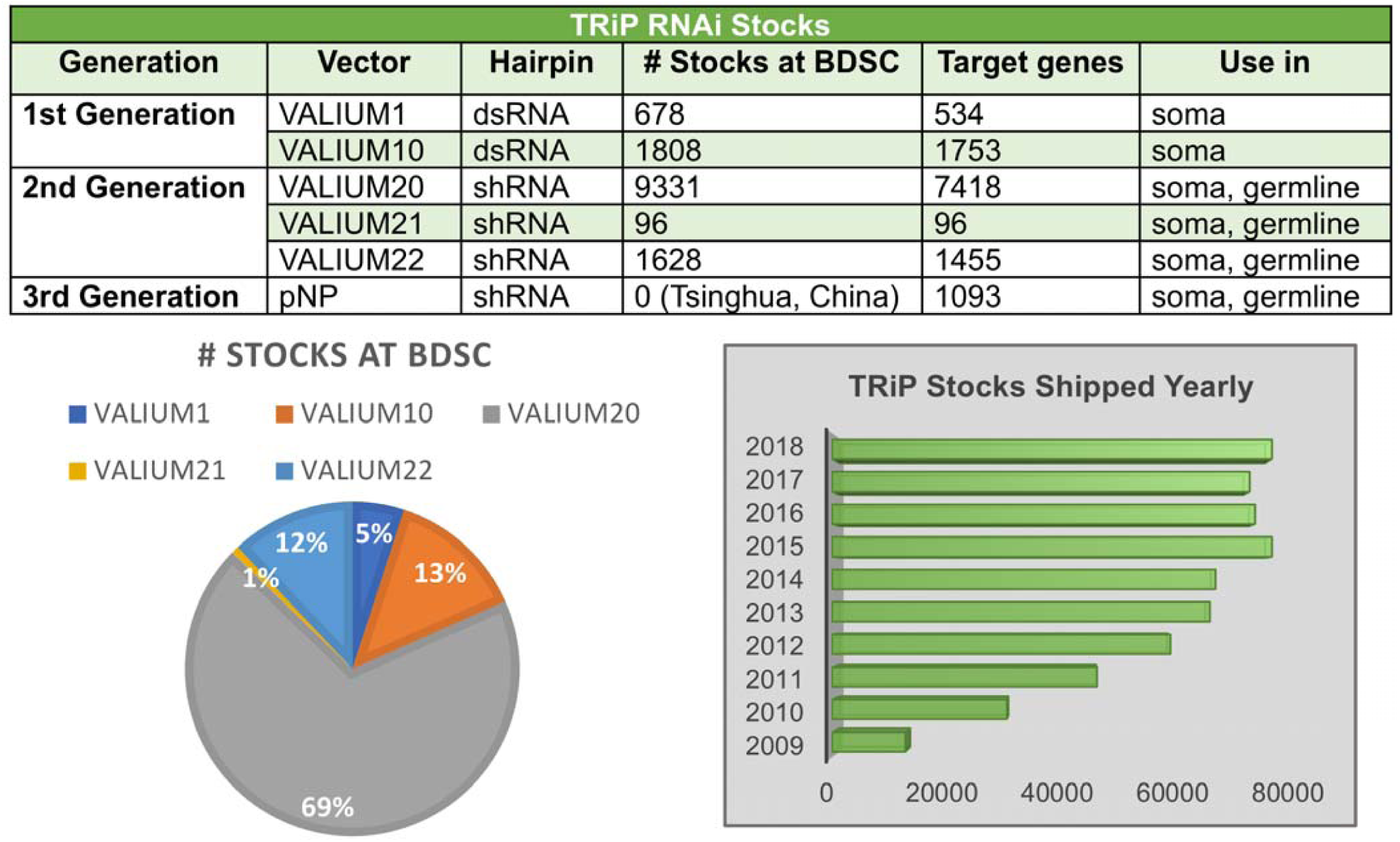
TRiP RNAi stocks and distribution. The first-generation VALIUM1 and VALIUM10 TRiP stocks contain long dsRNA hairpins. For VALIUM1, higher temperature fly culturing and UAS-Dicer 2 are best to achieve maximum knockdown. VALIUM10 was the best performing vector among our first series of related vectors, which were generated in an effort to optimize the various features of VALIUM1. The second-generation TRiP stocks contain short hairpins (shRNAs). VALIUM20, VALIUM21 and VALIUM22 have a modified scaffold derived from the microRNA miR-1 flanking the hairpin itself. VALIUM20 works well in the germline and is stronger than VALIUM10 in the soma. VALIUM22 (and VALIUM21) has features optimized for RNAi knockdown in the germline. Specifically, it has a P-transposase promoter instead of the *hsp70* basal promoter, and a K10 3’UTR instead of the SV40 3’UTR. Currently, BDSC is distributing ∼13,400 RNAi stocks, and to date have shipped over 700,000 TRiP stocks.

### TRiP-CRISPR collections

#### TRiP-CRISPR knockout (TRiP-KO)

CRISPR/Cas9 efficiently generates DSBs in *Drosophila* that can be used effectively to generate mutations or for genome engineering (Bassett et al. 2013; Gratz et al. 2013; Kondo and Ueda 2013; Ren et al. 2013; Yu et al. 2013; Sebo et al. 2014). Mutant animals or tissue-specific mosaics can be produced by simply crossing sgRNA-expressing flies to germline-specific Cas9 or somatic tissue-specific GAL4>Cas9 flies, respectively (Kondo and Ueda 2013; Port et al. 2014; Port and Bullock 2016). A number of recent studies in *Drosophila* have used transgenic CRISPR for LOF, for example to identify novel modifiers of tauopathies (Butzlaff et al. 2015; Dourlen et al. 2017) or Parkinson’s disease (Chen et al. 2017; Rousseaux et al. 2018); to model congenital disorders of glycosylation (Parkinson et al. 2016); to identify genes that regulate tumorigenesis (Das et al. 2013; Mishra-Gorur et al. 2019); and to study inflammation and immunity (Lee et al. 2018; Liu et al. 2018).

Recently, we and other groups (Meltzer et al. 2019; Port et al. 2019) started to generate libraries of transgenic sgRNA lines to complement existing RNAi collections. Unlike RNAi, which is often limited by incomplete penetrance due to residual gene expression (Echeverri and Perrimon 2006; Ma et al. 2006; Perkins et al. 2015), CRISPR/Cas9 targeted to the beginning of the open reading frame (ORF) of a protein-coding gene can generate frameshift mutations that strongly reduce or eliminate gene function. As with TRiP-RNAi lines, all TRiP-CRISPR stocks undergo rigorous quality control at the TRiP facility before being shipped to the BDSC for distribution. Available stocks are annotated on the DRSC/TRiP sgRNA Fly Stock Database (https://www.flyrnai.org/tools/grna_tracker/web/) and on FlyBase (Thurmond et al. 2019). As we build this new CRISPR collection, we encourage and receive gene target nominations from the community. As diagrammed in Fig. 2, TRiP-KO can be used for:

**Figure 2:**
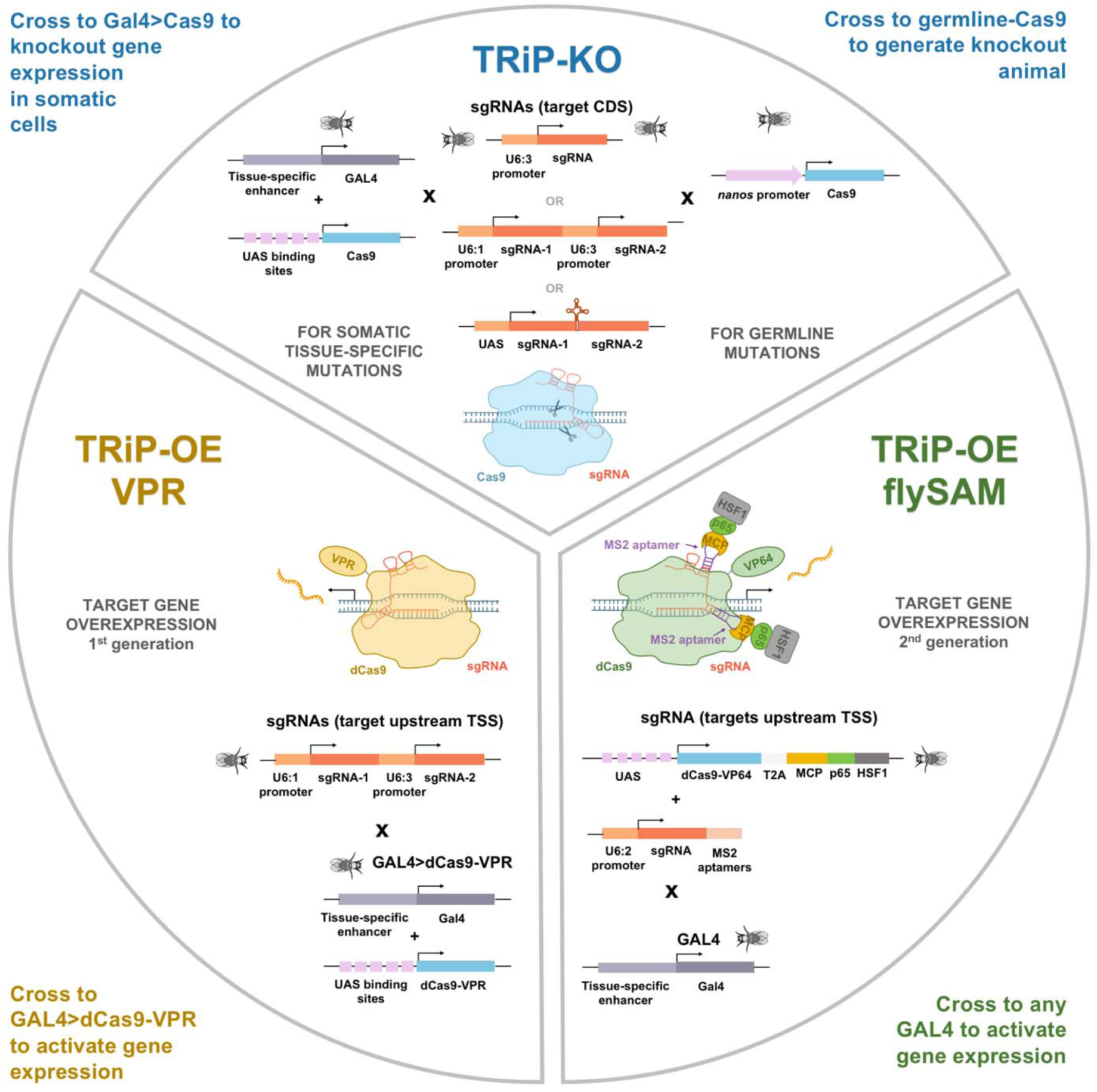
TRiP-CRISPR systems for *in vivo* gene knockout and overexpression. In the TRiP-CRISPR knockout collection (TRiP-KO), stocks express one or two sgRNAs targeting the coding sequence of a gene or genes. Crossing TRiP-KO stocks to a Gal4 line expressing Cas9 induces cleavage of the target site, non-homologous end joining (NHEJ) repair and insertion/deletion mutations (indels) in both germline and somatic tissue. Stocks in the TRiP-CRISPR overexpression (TRiP-OE) library express small guide RNAs (sgRNAs) targeting upstream of a gene transcription start site. Gene activation is triggered by co-expression of catalytically dead Cas9 (dCas9) fused to an activator domain, either VP64-p65-Rta (VPR) or Synergistic Activation Mediator (SAM).

1. **Germline KO by Cas9**. We produce fly stocks that constitutively express either one or two sgRNAs that target the coding sequence of a single gene. One or two sgRNAs per target are cloned into the pCFD3 and pCFD4 vectors, respectively (Port et al. 2014). To make a mutant fly strain, the sgRNA fly stock is crossed to flies with a germline-specific source of Cas9 (nanos-Cas9). CRISPR/Cas9-induced NHEJ occurs in the germ cells of the resulting progeny, which are then crossed to a balancer strain. The balanced flies are used to established stocks, which are screened for mutations by PCR.
2. **Somatic tissue mosaic KO by Cas9.** To generate mosaic tissues, an appropriate GAL4 line is used to drive expression of UAS-Cas9 in a specific cell population. When crossed to flies constitutively expressing one or more sgRNA, this results in mutations in the target gene only in the GAL4 expressing cells.

For all TRiP-CRISPR stocks, we used the DRSC Find CRISPR Tool (Ren et al. 2013) to pick the optimal sgRNA designs. With the KO collection, we selected the sgRNAs targeting the 1^st^ or 2^nd^ coding exons common to all isoforms with high efficiency scores and specificity scores. The efficiency scores we considered include the Housden efficiency score, which is based on a nucleotide position matrix (Housden et al. 2015), and the frameshift score, which predicts the likelihood of a frameshift happening based on microhomology sequences near the cutting site (Bae et al. 2014). At the same time, we also prioritize the designs with high specificity scores including the seed score (the length of unique seed region) and off-target score.

To date, we have produced 2070 TRiP-KO lines targeting community nominations, orthologs of human diseases, genes of unknown function, and autophagy-related genes, and other focused gene sets (Fig. 3). The first generation of TRiP-KO stocks have been made using the pCFD3 or pCFD4 vectors, in which the sgRNA is driven by the U6 promoter (Port et al. 2014). It was recently shown that in mosaic KO experiments combining tissue-specific-GAL4, UAS-Cas9 with U6 driven guides can result in some DSBs outside the GAL4 expression domain (Port and Bullock 2016). For this reason, TRiP-KO stock production has now shifted to using the pCFD6 vector (Port and Bullock 2016), in which sgRNA expression is under UAS control.

**Figure 3:**
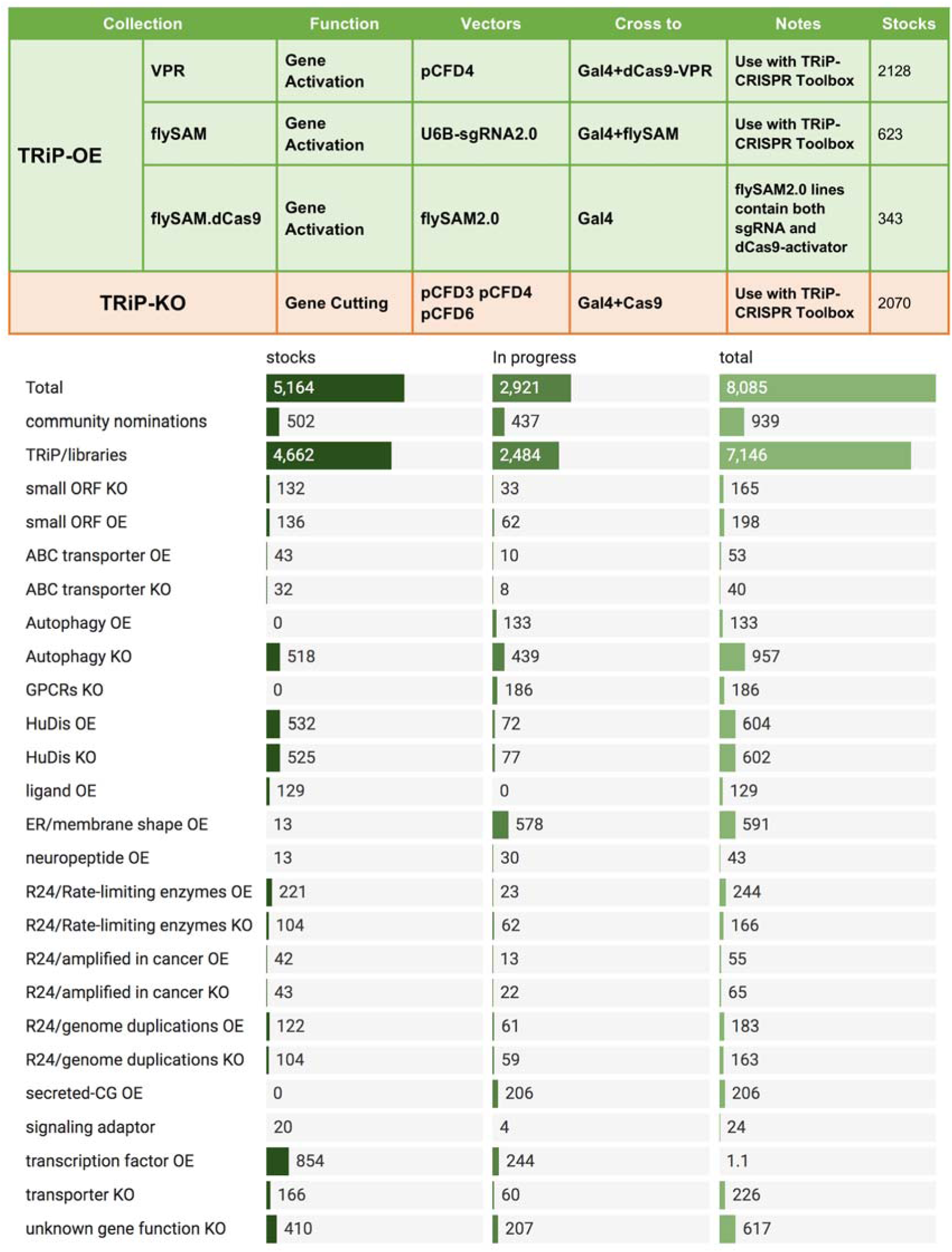
TRiP-CRISPR libraries. **(A)** There are three types of TRiP-OE sgRNA stocks: VPR, flySAM guide only, and flySAM.dCas9 with both guide and UAS-dCas9. TRiP-KO stocks are made in the pCFD series of vectors for either constitutive or UAS driven expression of single or double guides. **(B)** TRiP-CRISPR libraries in progress.

#### TRiP-CRISPR overexpression (TRiP-OE)

In contrast to the wealth of reagents available for LOF studies in *Drosophila*, the development of large-scale resources for GOF studies has lagged behind. This is a major gap in functional genomics, as mis- and overexpression screens provide an important complementary resource for elucidating gene function. Cas9 activators, in which dCas9 recruits transcriptional activation machinery to a DNA sequence upstream of the transcriptional start site (TSS) of a target gene, can potentially overcome these obstacles. Target specificity is conferred by 20 bp protospacer sequences in the sgRNA, such that production of reagents for CRISPR activators (CRISPRa) at genome-wide scale is feasible. We reported the first demonstration of effective CRISPR/Cas9-based transcriptional activation in flies (Lin et al. 2015), and have since developed and optimized two systems for CRISPRa *in vivo* (Chavez et al. 2016; Jia et al. 2018). Based on these methods, we have now generated a collection of transgenic TRiP-CRISPR overexpression (TRiP-OE) lines for gene activation (Fig. 2). TRiP-OE stocks express sgRNAs targeting genes upstream of the TSS. Gene activation is triggered by co-expression of catalytically dead dCas9 fused to an activator domain, either VP64-p65-Rta (VPR) or Synergistic Activation Mediator (SAM). sgRNAs for both systems are designed using the DRSC Find CRISPR Tool (Ren et al. 2013). We select sgRNAs targeting the region of 50 to 500 bp upstream the TSS of the targeted gene with high specificity scores.

With VPR, the TRiP-OE sgRNA stocks are crossed to a stock in which GAL4 directs expression of dCas9-VPR (Lin et al. 2015; Chavez et al. 2016). In the resulting progeny (*GAL4>dCas9-VPR; sgRNA-gene*), the gene of interest is overexpressed in the GAL4 domain. The flySAM method is based on the mammalian engineered protein complex (Konermann et al. 2015). In our version (Jia et al. 2018), a VP64 domain is fused to dCas9 and two additional activator domains, p65 and HSF1, are recruited to the complex via MS2 stem loops in the sgRNA tail. With flySAM, the first set of TRiP-OE sgRNA stocks were made in the U6B-sgRNA2.0 vector (Jia et al. 2018). To activate gene expression, these are crossed to a stock in which GAL4 directs expression of UAS-flySAM. In the more recent flySAM.dCas9 collection, the stocks are made with the flySAM2.0 vector, which contains both the flySAM protein complex and the sgRNAs, thus gene activation is achieved by simply crossing to the GAL4 line of interest. The flySAM method gives considerably greater levels of activation compared to VPR and is comparable to the UAS overexpression system.

Importantly, the CRISPRa system has several advantages over UAS-cDNA overexpression as it allows: (1) overexpression of untagged wildtype gene products; (2) overexpression of all of the relevant splice isoforms for a given cell-type; (3) preservation of the 3’UTR, which is important as it contains regulatory information such as miRNA binding sites; and (4) overexpression of ORFs that are long, contain repeat sequences, or are otherwise intractable using a cDNA approach due to cloning difficulties. Finally, generating a useful genome-wide collection requires that the large majority of the lines generated are functional. To estimate the proportion of TRiP-OE sgRNA lines that function as predicted, we previously created a panel of transgenic fly stocks expressing sgRNAs and analyzed activation using qPCR (Ewen-Campen et al. 2017). Seventy-five percent of these sgRNA transgenes led to a greater than three-fold increase in transcript levels when combined with dCas9-VPR, and 58% caused a greater than eight-fold increase, a success rate comparable to the proportion of transgenes in current RNAi stock collections that confer effective knockdown. In addition, several of the lines that do not appear up-regulated when tested via qPCR do in fact produce specific phenotypes, indicating that these success rates are in fact likely to be underestimated.

We also tested the stringency of the system with respect to off-target gene activation. This is a concern, since due to the compactness of the *Drosophila* genome, multiple genes could be within range of activation from a single sgRNA. Using qPCR, sgRNAs were analyzed *in vivo* for transcriptional activation of the target gene as well as neighboring genes within a ∼20 kb window. For four our of five target genes, there was no off-target activation (Fig. 4A). Next, we tested TRiP-OE induced protein expression of a gene containing a GFP protein trap (Fig. 4B). We combined the wing pouch-specific nub-GAL4 driver with UAS-dCas9-VPR, then crossed to the TRiP-OE guide stock targeting CG6767 (GS01642), the GFP MiMIC protein trap in CG6767 (MI09551-GFSTF.2), or both together. When all components were expressed, we observed strong TRiP-OE induced expression of the GFP-tagged CG6767 protein. Importantly, GFP upregulation was strictly confined to the nub-GAL4 domain in the wing pouch, indicating that the CRISPRa system allows tight spatial control of gene expression. Finally, we tested the flySAM system for toxicity. Previously, it had not been possible to apply SAM *in vivo* in flies because ubiquitous expression of UAS:MCP-p65-HSF1 is lethal in the absence of any sgRNA (Ewen-Campen et al. 2017). We therefore tested whether flySAM can be expressed *in vivo* without toxicity by crossing to a panel of GAL4 driver lines with or without control sgRNA (Fig. 4C). We observed normal survival rates and no visible phenotypes for most tissue-specific GAL4 lines. However, we did observe complete lethality with one ubiquitous driver (tubulin-GAL4), and we observed moderate lethality and phenotypes with some strong tissue-specific drivers (eg. Mef2-GAL4 and ey-GAL4). These results suggest that very strong GAL4 drivers should be avoided for flySAM overexpression experiments.

**Figure 4:**
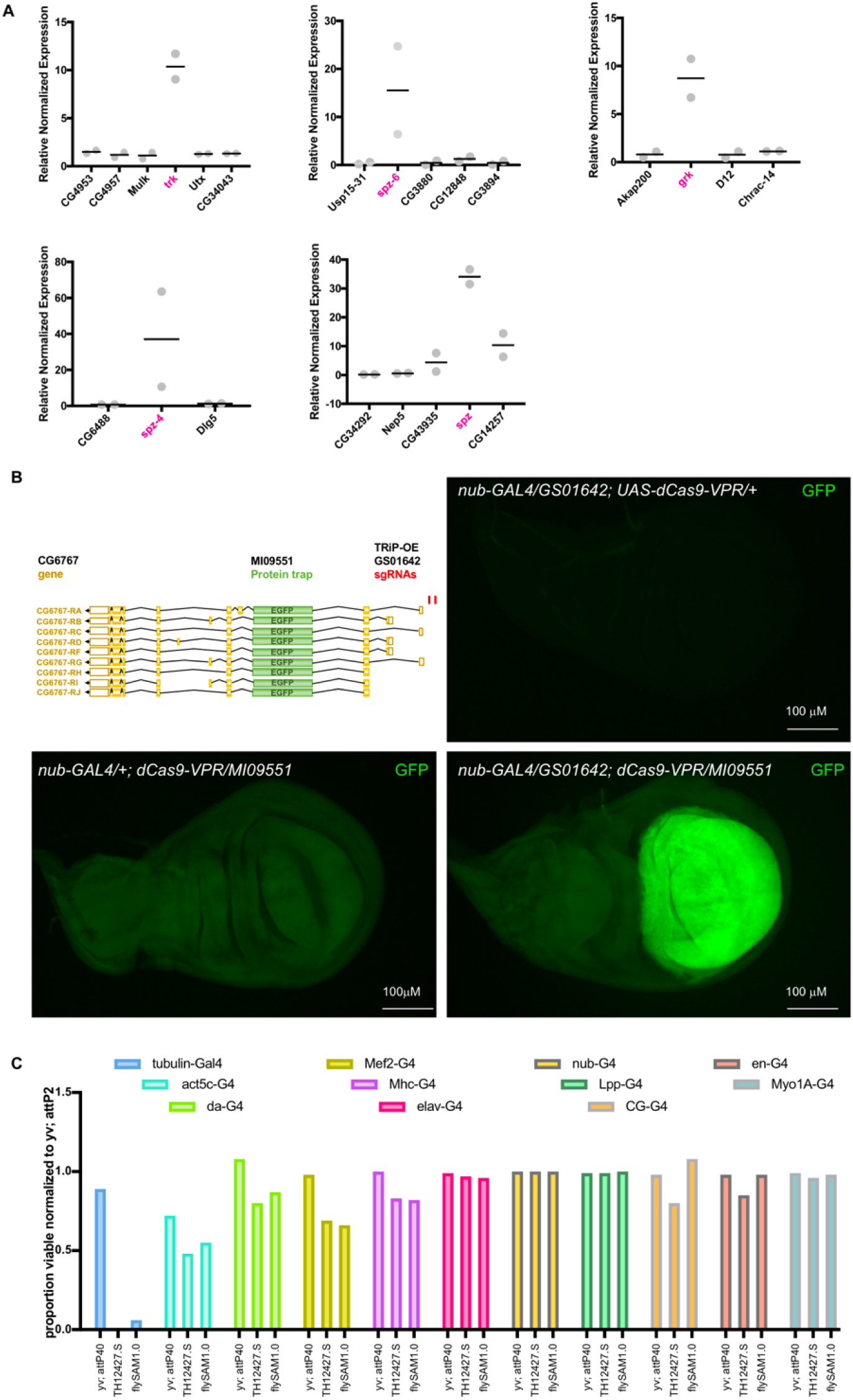
TRiP-OE systems provide robust on-target gene activation. **(A**) Larvae of the genotype *tubulin-GAL4 > UAS-dCas9-VPR, sgRNA* were analyzed for transcriptional activation of target gene as well as neighboring genes within a window of ∼20 kb. With one exception (*spz*), there was no off-target activation. Target genes are indicated in magenta. **(B)** TRiP-OE activation of tagged endogenous protein. Larvae of the genotype *nub-GAL4 > UAS-dCas9-VPR* were crossed to TRiP-OE sgRNA GS01642 (targeting CG6767), a GFP protein trap in CG6767 (MI09551-GFSTF.2), or both. Strong TRiP-OE induced expression of the GFP-tagged CG6767 protein was observed in the nub-GAL4 domain. **(C)** A panel of GAL4 drivers were crossed to controls (*yv; attp40* and *yv; attP2*), UAS-flySAM2.0 with non-targeting sgRNA (TH12427.S), or UAS-flySAM1.0 (no guide). The proportion of viable offspring were recorded and normalized to the *yv; attP2* control. Normal survival rates and no visible phenotypes for most tissue-specific GAL4 lines. Complete lethality was observed with one ubiquitous driver (tubulin-GAL4) and moderate lethality with some strong tissue-specific drivers (e.g. Mef2-GAL4).

To date, we have produced 3094 TRiP-overexpression (TRiP-OE) lines targeting transcription factors, small ORFs, orthologs of human diseases, kinases, autophagy-related genes, and other focused libraries (Fig. 3, Fig. S1). The majority of TRiP-OE flies were made using the VPR system. However, due to the greater levels of gene activation, as well as the ease of use, TRiP-OE stock production has shifted to using the flySAM system exclusively.

### Toolbox

In addition to the RNAi stocks, the TRiP, via the BDSC, also provides a “TRiP Toolbox” stock collection (Table S2) that includes injection stocks for labs wishing to generate their own RNAi lines and commonly used GAL4 lines with UAS-Dcr2, which is useful to enhance message knockdown when combined with long dsRNAs but not shRNAs. Because TRiP-KO and TRiP-OE VPR lines contain only the sgRNA, it is necessary to express Cas9 separately in the tissue of interest. For this reason, we have also produced a TRiP CRISPR/Cas9 toolbox set of GAL4/GAL80ts/UAS stocks that allow spatial and temporal expression of Cas9 or dCas9-VPR (Table S2). A total of 55 TRiP CRISPR/Cas9 toolbox lines are complete and have been shipped to BDSC for distribution.

### Human disease-related orthologs RNAi reagent library

To facilitate the use of *Drosophila* to study human diseases, we assembled and characterized a resource, which we refer to as the Human Disease-TRiP (HuDis-TRiP) resource, consisting of RNAi fly stocks targeting *Drosophila* orthologs of genes implicated in human disease. This resource will enable the biomedical research community to quickly determine if a *Drosophila* model is relevant to their studies and if so, allow them to embark on detailed follow-up analyses.

To identify the set of fly genes orthologous to genes known or suspected to be associated with human diseases, we selected 4,850 human disease genes described in the Online Mendelian Inheritance in Man (OMIM) database (Amberger et al. 2015; Amberger et al. 2019) as “with known sequence and phenotype” (omim.org/statistics/geneMap). We then mapped the human genes to fly genes using version 3 of our DIOPT tool (Hu et al., 2011) and selected those with DIOPT scores >=2 (best score, forward or reverse search) or DIOPT score >=4, identifying 3,017 genes (10 was the max score for DIOPT vs3). In an analysis of these genes in the literature, we found that as a group, they are associated with a large number of top-level medical subject heading (MeSH) terms relevant to diseases of interest, e.g. nervous system diseases, musculoskeletal diseases, eye diseases, skin diseases, digestive system diseases, cardiovascular diseases, neoplasms, metabolic diseases, etc. (Fig. 5). From this set of ∼3000 genes, we also extracted a set of 670 high-confidence *Drosophila* orthologs of high-confidence disease-associated human genes using DIOPT vs3 to map OMIM human disease genes to fly with very strong DIOPT score filter of 8 or higher. To date we have produced a total of 2,487 shRNA stocks targeting the full list of 3,017 genes (Fig. 5) with >95% coverage of the 670 high confidence set. A sortable list of HuDis stocks is available at https://fgr.hms.harvard.edu/hudis-trip-fly-stocks. The HuDis website contains links to the human genes at NCBI, the fly genes at FlyBase, DIOPT results (linked to from orthology scores), BDSC stock IDs, and RNAi screen data in RSVP Plus. The information on the website is meant as a high-confidence starting off point; it is not a comprehensive list of fly gene-human disease gene relationships.

**Figure 5:**
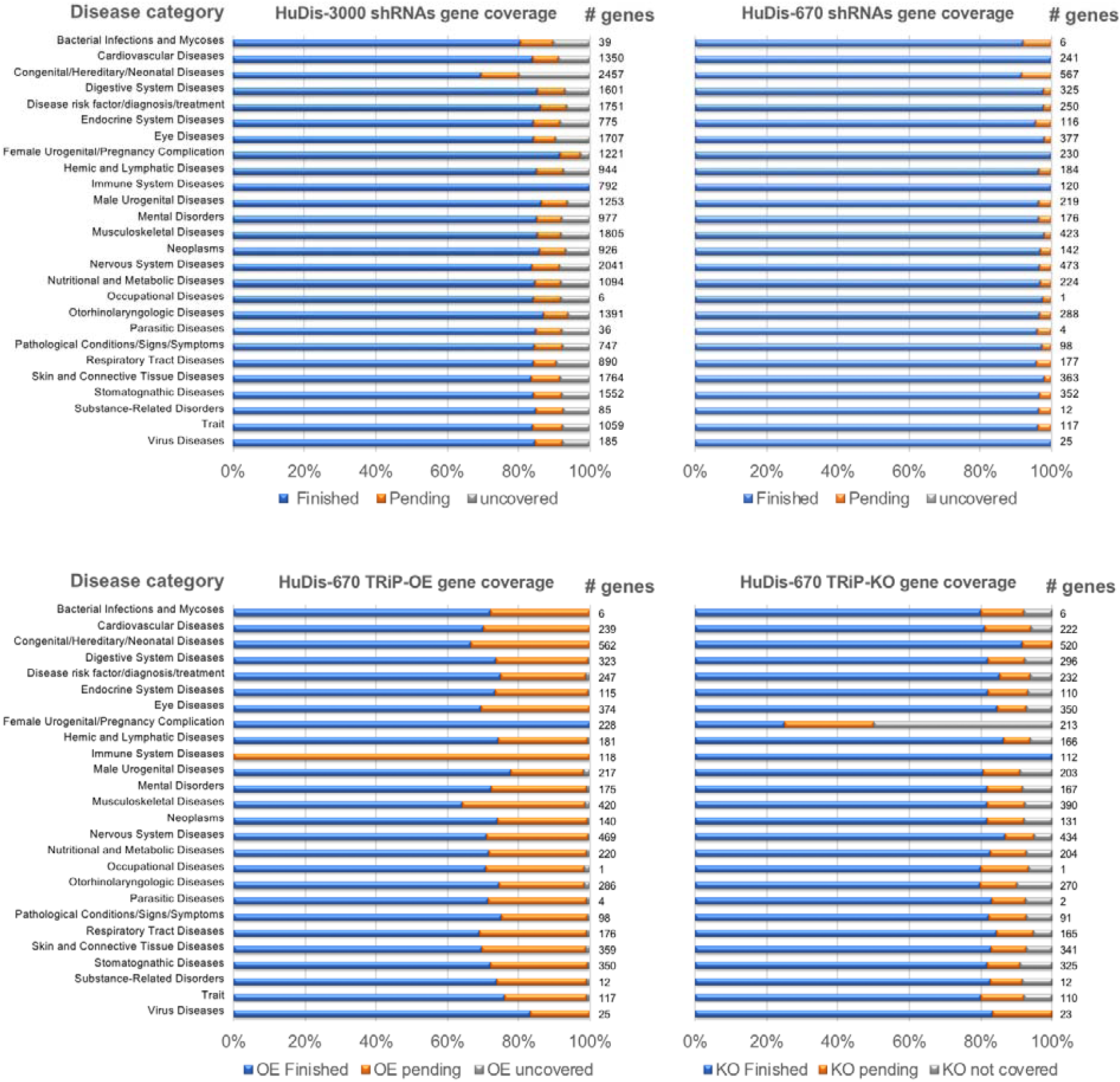
Summary of shRNA and sgRNA production for human disease orthologs. Production status of shRNA and sgRNA for human disease gene ortholog reagent libraries, organized by top-level medical subject heading (MeSH) terms. HuDis-3000 is the full set of ‘HuDis’ gene set orthologs with DIOPT vs.3 scores >=2 (best score, forward or reverse search) or DIOPT vs.3 score >=4. HuDis-670 is a smaller set of high confidence orthologs with DIOPT vs.3 scores >=8.

To validate the HuDis RNAi stocks, we first performed an initial characterization of ∼1,200 RNAi lines targeting the high-confidence set of 670 genes using two GAL4 driver lines: actin5C-GAL4 and daughterless-GAL4, which are expressed in most or all cells and stages, identifying RNAi knockdown that results in lethality. As shown in our workflow diagram (Fig. 6A), we then crossed the RNAi lines to a panel of selected GAL4 drivers to generate phenotypes in specific body parts (muscle and nervous systems, developing adult eye, wing, thorax, etc.) and at specific developmental stages. Flies were examined for visible morphological adult phenotypes, male and female sterility, and locomotor defects. We also measured sugar and triglyceride levels to determine whether the overexpressed gene affects overall metabolism. The full list of morphological and metabolic phenotypes is listed in Table S3. Based on these experiments, we identified 213 genes that were lethal when knocked down by ubiquitous-GAL4 with at least two RNAi lines (Fig. 6B). Of these, 163 gave the same phenotype with multiple RNAi lines in the follow-up studies with tissue specific GAL4s (Table S3).

**Figure 6:**
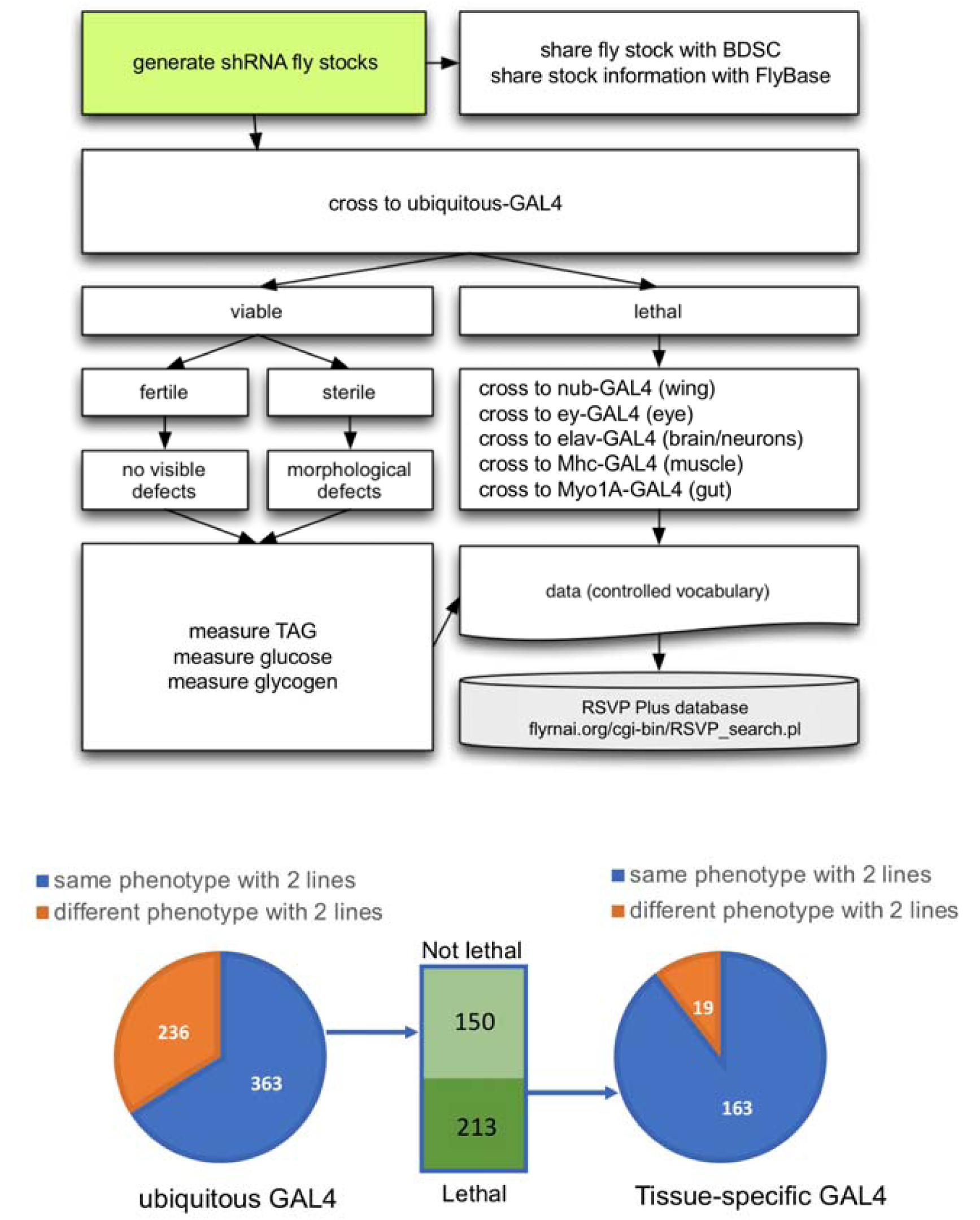
Characterization of HuDis RNAi lines. **(A)** Workflow diagram. We defined a set of high-confidence orthologs of high-confidence human disease-associated genes and generated a corresponding set of shRNA fly stocks. All available RNAi lines targeting the gene set (∼1200 fly stocks) were screened with ubiquitous GAL4 drivers (act5c and daughterless). Fly stocks were subcategorized as lethal or viable. The former were tested for sterility, gross morphological phenotypes, and metabolic defects. The latter were tested with a panel of tissue-specific GAL4 drivers. All phenotypic data was uploaded to RSVP Plus. **(B)** For 363 of ∼600 genes, results with two RNAi reagents were concordant, either both lethal or both viable. Knockdown of 213 of the genes resulted in lethality and 163 of these gave concordant phenotypes with multiple RNAi lines using a panel of tissue-specific GAL4 drivers.

HuDis-TRiP stocks were made in the VALIUM20 vector. Typically, ∼15% of shRNA stocks made in VALIUM20 do not homozygose due to leaky expression of the transgene (Qiao et al. 2018). Therefore, we also generated a resource of 1000 lines targeting human disease orthologs using the pNP vector (Qiao et al. 2018). The pNP stocks have reduced basal shRNA expression and RNAi specificity compared to VALIUM series stocks and augment the gene coverage of the collection.

Altogether, the HuDis-TRiP fly stock collection provides a resource for understanding the role of the human disease gene orthologs and for modeling human diseases in flies.

### Human disease-releated orthologs reagent CRISPR library

In addition to the HuDis-TRiP RNAi library, we have also assembled TRiP-KO and TRiP-OE libraries targeting the high-confidence HuDis orthologs, as well as putative rate-limiting enzymes, genes that are overexpressed in cancer, and genes mapped to microduplication syndromes (Table S4). To date, we have generated TRiP-OE lines covering 74% and 44%, respectively, of the high confidence and complete sets of HuDis orthologs for which we were able to design sgRNAs (Fig. 5, Fig. S1). We have likewise generated TRiP-KO lines covering 76% and 45%, respectively, of the high confidence and complete sets of HuDis orthologs for which we were able to design sgRNAs (Fig. 5, Fig. S1). The TRiP-KO stocks serve as a complement to RNAi lines for LOF studies and the TRiP-OE stocks serve as a means to test overexpression models, as might be particularly relevant for human diseases associated with dominant mutations, polynucleotide expansions, translocations, amplification, or duplication syndromes. As noted previously, GOF analysis can also help elucidate gene function when LOF produces no phenotype. As an example, the *suppressor of rudimentary gene [su(r)]*, encodes dihydropyrimidine dehydrogenase, which catalyzes the rate-limiting step of pyrimidine degradation. Human dihydropyrimidine dehydrogenase (DPYD) expression is positively associated with aggressive tumor characteristics in several cancers and is predictive for poor patient prognosis (Mizutani et al. 2001; Horiguchi et al. 2002; Terashima et al. 2003; Zhu et al. 2018). In *Drosophila*, classical LOF *su(r)* alleles or RNAi targeting this gene produce no visible phenotypes. Rather, the function of *su(r)* in pyrimidine metabolism was only identified by suppression of the truncated wing phenotype exhibited by *rudimentary (r)* mutants in which de novo pyrimidine biosynthesis is blocked (Norby 1970; Rawls and Porter 1979). Using the wing-specific nubbin-GAL4 driver combined with UAS-dCas9-VPR and a TRiP-OE HuDis stock targeting *su(r)*, we were able to generate a strong ‘rudimentary-like’ small wing phenotype (Fig. 7).

**Figure 7:**
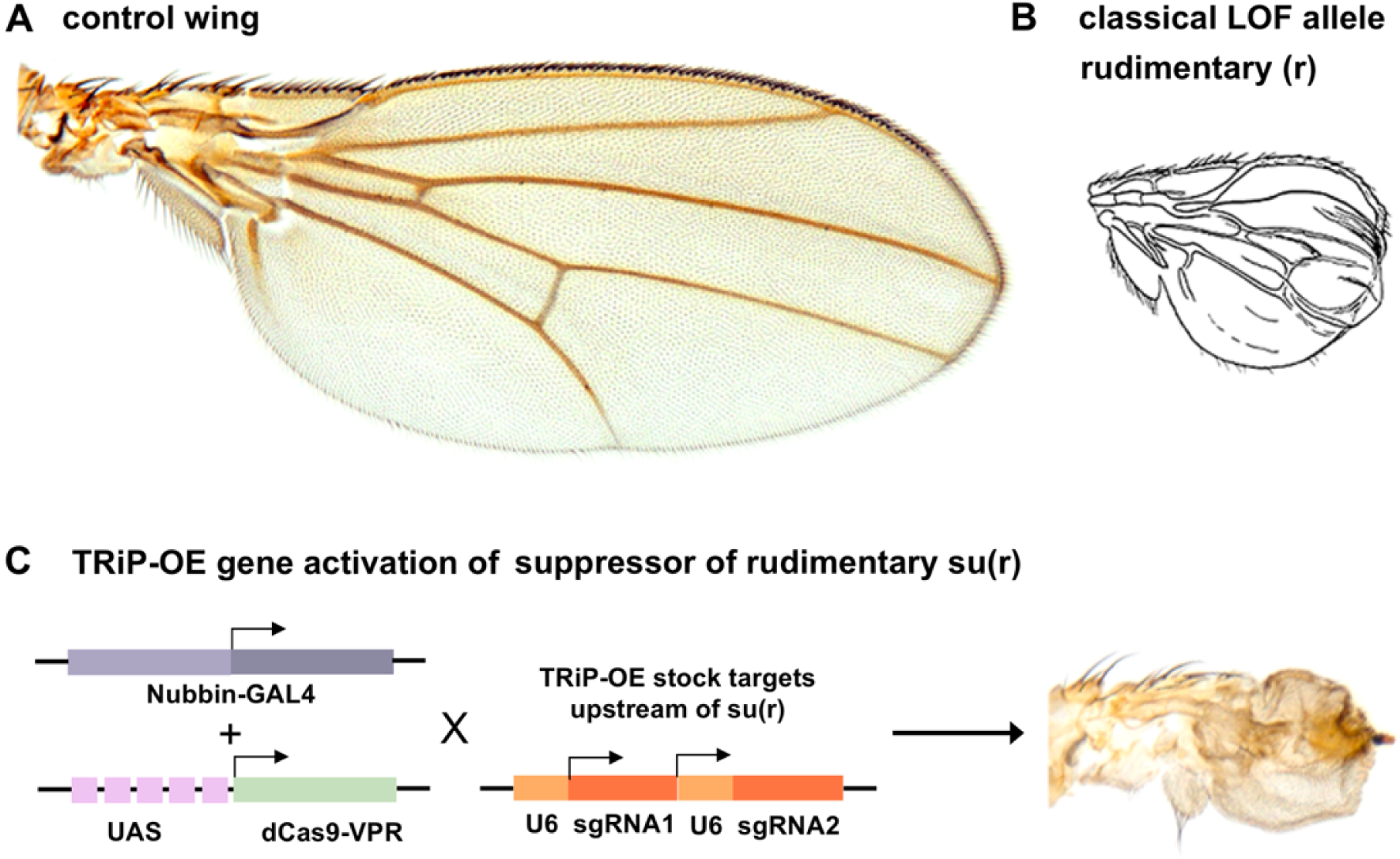
TRiP-OE model for *dihydropyrimidine dehydrogenase* overexpression. **(A)** Control fly wing**. (B)** The *rudimentary (r)* gene encodes the *Drosophila* carbamoyl-phosphate synthetase 2 (CAD), which catalyzes the rate-limiting step of pyrimidine synthesis. Classical *r* alleles cause dramatic wing tissue undergrowth, as shown in a line drawing from the ‘Red Book’ (Lindsley et al. 1992). **(C)** nubbin-GAL4 driving UAS-dCas9-VPR + TRiP-OE sgRNA targeting suppressor of rudimentary *[su(r)]*, which encodes dihydropyrimidine dehydrogenase, an enzyme that catalyzes the rate-limiting step of pyrimidine degradation. As expected, overexpression of *su(r)* phenocopies the LOF of *r* phenotype.

### TRiP data access and curation

Information about TRiP plasmid vectors, stocks and/or protocols are available (1) in published papers (Ni et al., 2008, 2009, 2011; Perkins et al. 2015), (2) at our website, viewable online (https://fgr.hms.harvard.edu/trip-in-vivo-fly-rnai), (3) for RNAi reagents, at our website, as a downloadable Excel file of TRiP fly stocks (see https://fgr.hms.harvard.edu/trip-in-vivo-fly-rnai), (4) at our website, searchable using Gene Lookup (https://www.flyrnai.org/cgi-bin/DRSC_gene_lookup.pl), Updated Targets of RNAi Reagents (UP-TORR) (https://www.flyrnai.org/up-torr/), or RSVP (see below), (5) for CRISPR sgRNA fly stocks, using or our sgRNA nomination and production tracking database (https://www.flyrnai.org/tools/grna_tracker/web/), (6) at FlyBase, and (7) at the BDSC. In addition, validation and phenotype data for RNAi and CRISPR can be accessed through the RSVP Plus online database (https://www.flyrnai.org/cgi-bin/RSVP_search.pl) (Fig. 8). RSVP Plus is an expanded version of our RSVP database (Perkins et al. 2015), modified to accommodate CRISPR data as well as RNAi data. We populate RSVP Plus in the following ways: (1) with unpublished data collected in-house or other research groups, (2) with published data shared in bulk file formats (e.g. Excel spreadsheets) by other research groups, (3) through community input online, and (4) through capture of RNAi or CRISPR fly stock phenotype data curated by FlyBase. Thus, RSVP Plus, which we maintain and regularly update, allows users to search and view information about knockdown, knockout, or over-expression phenotypes, including qPCR data and phenotypes (text or images) for specific reagent-GAL4 combinations. RSVP Plus also includes a modified user interface that lets external users upload data. At RSVP Plus there are currently >11,000 data entries for about 5,500 TRiP lines representing about 3,900 fly genes. In addition, RSVP Plus contains 23,451 data entries from FlyBase for 17,782 RNAi lines representing 11,346 genes, including lines from the Vienna Drosophila Resource Center (VDRC) and the National Institute of Genetics-Japan (NiG-Fly). Altogether, researchers will be able to rapidly identify and make use of this data resource to identify effective reagents and design efficient experiments based on published and new data.

**Figure 8:**
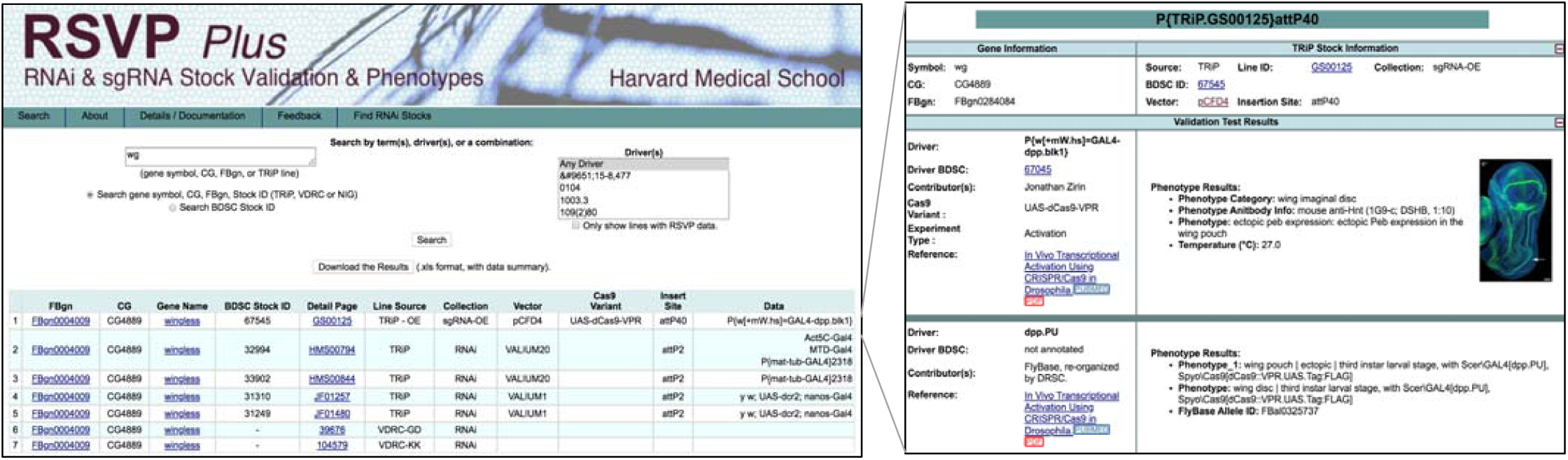
RSVP Plus database of RNAi and CRISPR phenotypes. Shown here are the Search and Details pages for a search with the *wingless* (*wg*) gene. RSVP Plus allows users to view information about knockdown, knockout, and overexpression efficiency (qPCR data) and phenotypes (text and when available, images) for specific combinations of RNAi or sgRNA fly stocks with Gal4 drivers and when relevant, Cas9 versions (e.g. Cas9 or dCas9-VPR or flySAM). RSVP Plus includes results curated by FlyBase for TRiP reagents or other major stock collections (e.g. VDRC, <u>NIG-fly Japan</u>).

### Concluding Remarks

The TRiP platform has been used to generate more than 20,000 RNAi and CRISPR fly stocks for the research community. These resources, distributed to the community by the BDSC, provide powerful, versatile and transformative tools for gene knockdown, knockout, activation, and genome engineering. Researchers can now easily access fly stocks useful to ‘dial down’ or ‘dial up’ genes covered by the collection, in a given stage or tissue (made possible with the large collection of available Gal4 drivers), and as a result, facilitate a near limitless array of single-gene and multiplexed genetic experiments, as well as facilitate further analysis and integration of the resulting phenotypic data.

## Acknowledgements

We thank Filip Port for transgenic Cas9 stocks and cloning advice. We thank the Bloomington Drosophila Stock Center and FlyBase for their support on TRiP fly stock data and resource deposition, as well as for their long-term and continued support of fly stock and data distribution to the *Drosophila* community. We thank the Harvard Medical School (HMS) Research Computing group for support relevant to our bioinformatics workflows and online resources. We thank the HMS Image Data Management Core and HMS Biopolymers Facility for relevant services and support. We also thank the Dana Farber/Harvard Cancer Center (DF/HCC) in Boston, MA, for use of the DNA Resource Core Facility. The DF/HCC is supported in part by NCI Cancer Center Support Grant NIH 5 P30 CA06516. The TRiP at HMS is supported by NIGMS R01 GM084947, with additional support from NCRR/ORIP R24 RR032668, NCRR/ORIP R24 OD021997, and NIGMS P41 GM132087. SEM is also supported in part by the DF/HCC and NP is an investigator of the Howard Hughes Medical Institute.

## Supplementary Methods

### The TRiP RNAi collection

#### RNAi vectors

The TRiP has generated a series of 22 knockdown vectors, the VALIUM series vectors (for Vermilion-AttB-Loxp-Intron-UAS-MCS), to facilitate incorporation of RNAi hairpins into attP landing sites (Ni et al. 2008; Ni et al. 2009; Ni et al. 2011; Perkins et al. 2015). All VALIUM vectors contain a wild type copy of *vermilion* as a selectable marker and an attB sequence to allow for phiC31 targeted integration at genomic attP landing sites (Groth et al. 2004). The VALIUM vectors were also designed with two pentamers of UAS sequences, one of which can be removed using the Cre/loxP system. In addition to manipulating the number of UAS sequences, the level of RNAi knockdown can also be altered by using Gal4 lines of various strengths, rearing flies at different temperatures, or via co-expression of UAS-Dicer2 (Dietzl et al. 2007) in the case of long double stranded RNAs (dsRNAs). The first-generation knockdown vectors chosen by the TRiP for RNAi stock production were VALIUM1 and VALIUM10. Both allow expression of long dsRNA hairpins, usually between 400 and 600 bp. These are very effective for RNAi in somatic tissues but are not as effective in the female germline (Ni et al., 2008; Ni et al., 2009). Subsequently, we showed that shRNAs containing a 21 bp targeting sequence embedded into a micro-RNA (miR-1) backbone are very effective for gene knockdown in both the germline and soma. For shRNA expression we developed the second-generation knockdown vectors, VALIUM20, VALIUM21 and VALIUM22 (Ni et al. 2011). Subsequent TRiP lines were generated with shRNAs in VALIUM20 (for knockdown in germline or soma) or VALIUM22 (mostly germline). Finally, since some researchers prefer to use *mini-white* as the selectable marker for transgenesis, we also generated new versions of the VALIUM vectors in which *vermilion* is replaced with *white* (WALIUM10, WALIUM20 and WALIUM22). Further information about the TRiP vectors in available at the TRiP website (https://fgr.hms.harvard.edu/trip-plasmid-vector-sets).The vectors are distributed by the <u>Drosophila Genome Resource Center</u> (https://dgrc.bio.indiana.edu/).

#### Generation of the collection

Our pipeline is as follows: (1) design shRNAs targeting each gene, (2) order oligos corresponding to shRNA sequences, (3) clone them into the expression vector and sequence verify, (4) inject into embryos, (5) isolate and balance transgenic flies, (6) sequence verify the transgenic stocks, (7) add stock information to our in-house database and coordinate with Flybase, (8) ship to BDSC for distribution.

To design an shRNA for a given gene, we first determine the sequences of all exons (or when appropriate, the sequences of exons common to all transcripts), then determine all possible 21 bp sub-sequences of the sense or antisense strands, and finally, compare these to all genes. Sub-sequences with 16 bp or longer stretches of identical nucleotide matches to other genes are removed from consideration. Each subsequence is assigned a score based on a formula by Vert et al. (Vert et al. 2006) and the top scoring sub-sequences are selected.

All fly stocks generated at the TRiP are inserted into one of two attP sites, attP40 on the left arm of the second chromosome at 25C6 or attP2 on the left arm of the third chromosome at 68A4. These sites were selected for their abilities to provide high levels of induced expression of the transgenes but maintain low basal expression when the transgenes are not induced (Markstein et al. 2008). The landing site chosen by the TRiP for hairpin insertion is guided first by the preference of the community member nominating the gene and second by the TRiP. If a TRiP stock for a particular gene is available in one location, a second TRiP stock for the same gene will be generated in the other location. shRNA constructs are generated individually in 96-well plates, or selected from shRNA libraries generated in VALIUM20 and VALIUM22 starting from a pool of 83,256 unique shRNA oligonucleotides synthesized on glass slide microarrays (Ni et al. 2011). Early in the project, constructs were injected individually into attP40- or attP2-bearing lines and transformants were recovered. As only one attB insert can integrate by phiC31-mediated recombination into an attP site (Groth et al. 2004), we later injected pools of constructs, established transgenic lines, and then subsequently characterized the inserted DNA by Sanger sequencing.

We produce the lines in the Perrimon lab, with injection companies such as Bestgene, or with the help of two outside groups, the National Institute of Genetics (NIG) in Japan (coordinated by Drs. Shu Kondo and Ryu Ueda) and the THFC at Tsinghua University in China (coordinated by Dr. Jianquan Ni). Importantly, these outside labs use established TRiP nomenclature and send the lines they generate to the TRiP at HMS, where they are checked for quality. All completed stocks are annotated on the TRiP website (http://fgr.hms.harvard.edu/) and on FlyBase, then transferred as soon as possible to the BDSC for distribution to the community. Select stocks are also available from the NIG and the THFC.

#### qPCR primers

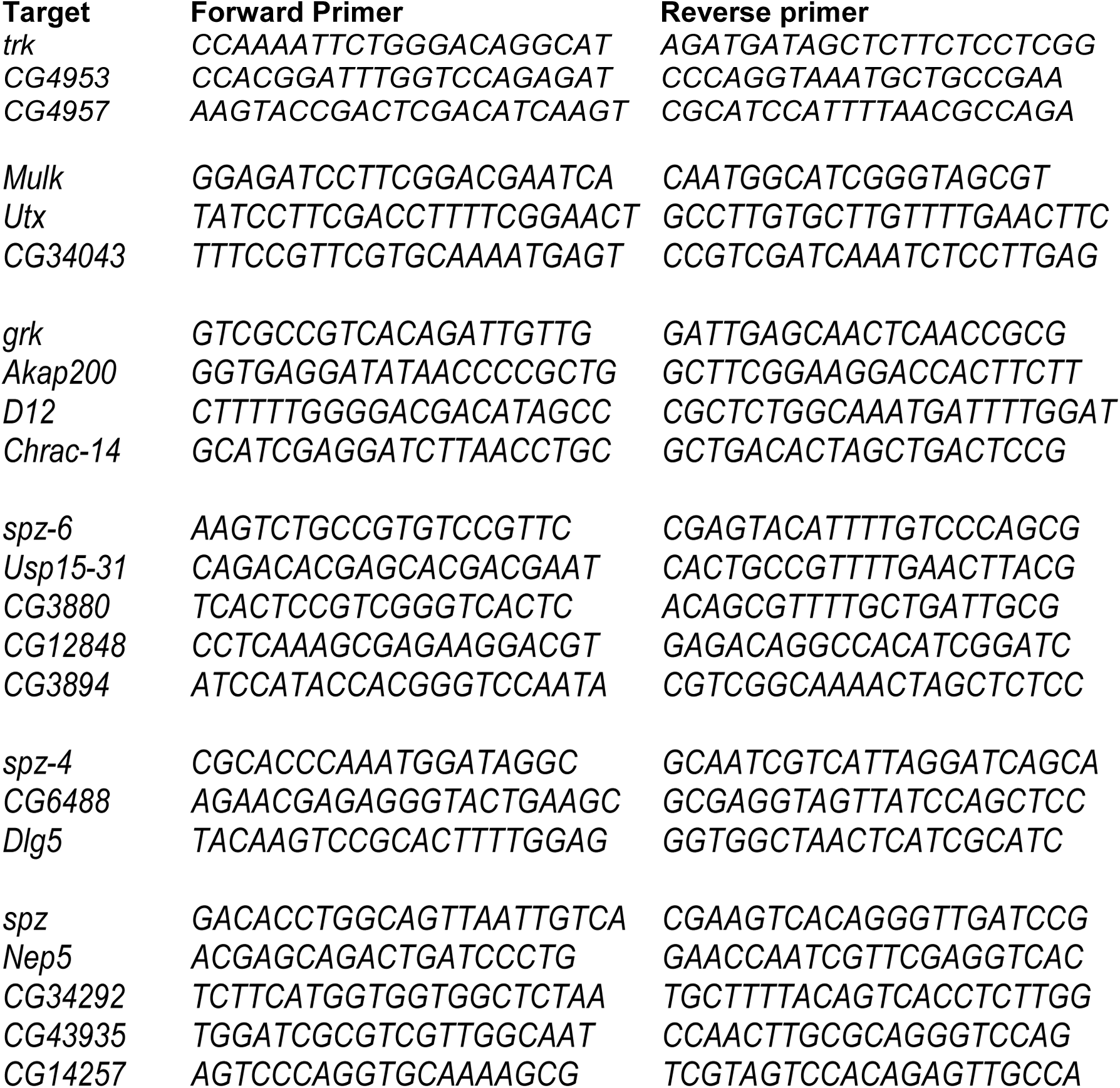

**Figure S1:**
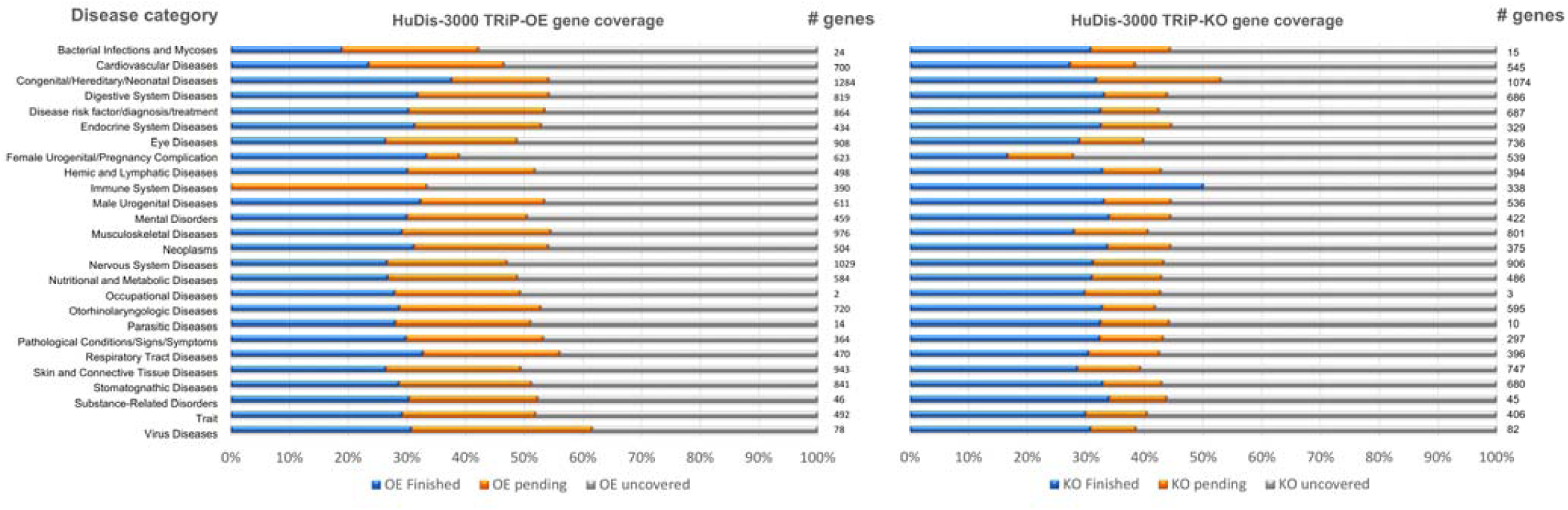
Summary of sgRNA production for human disease orthologs (HuDis-3000)

